# Validated microsatellite markers for *Gyrodactylus salaris*: a toolkit for individual identification and genetic studies

**DOI:** 10.64898/2026.04.22.719836

**Authors:** Heidi Aisala, Haakon Hansen, Jaakko Lumme

## Abstract

Microsatellite markers remain essential for individual-level genetic work in taxa where genome-wide methods are not yet routinely feasible due to extremely low DNA yields per specimen. In *Gyrodactylus*, even the most recent reference genomes have required pooling thousands of individuals, leaving a practical gap between genome-scale resources and individual-level analyses. Here we present a genome-informed microsatellite panel, developed by selecting single-copy loci with non-repetitive flanking regions and assembling all markers into a single multiplex PCR. Marker identity and performance were verified via amplification tests, Sanger sequencing, and cross-laboratory genotyping, confirming that the same samples generated identical fragment-size profiles in both laboratories. Long tandem repeats occasionally prevented exact repeat-count determination, yet allele-size classes were discrete and reproducible across replicates. The panel enables rapid individual identification and reliable strain and lineage assignment. It also offers a practical starting point for population-genetic and evolutionary studies that require individual-level data.

## Introduction

Genome-wide methods such as whole-genome sequencing/resequencing, reduced-representation libraries (e.g. RAD/GBS), and large SNP panels are now widely used in population and conservation genetics (Allendorf et al. 2010, Andrews et al. 2016, Hohenlohe et al. 2021, Luikart et al. 2003). However, these approaches depend on high-quality DNA and individual-level sequencing libraries, which are difficult to obtain from very small organisms with extremely low per-individual DNA yields (Andrews et al. 2016, Hodel et al. 2016). As a result, earlier genetic studies in such systems have often relied on conserved rDNA or mitochondrial markers, which lack the variation needed to differentiate individuals and closely related strains (Hodel et al. 2016). In our focal genus *Gyrodactylus*, these markers have been used extensively, and their limitations are well established (Hansen et al. 2007). Beyond these, genome-wide work in *Gyrodactylus* has generally required large numbers of individuals per sample, including genome-wide resequencing projects, all published nuclear genomes, and efforts to develop more variable molecular markers (Faria et al. 2011, Hahn et al. 2014, Konczal et al. 2020a, Konczal et al. 2020b, Zhang et al. 2025). Together, these studies show that producing reliable data at the level of single worms has remained technically challenging in this group, which continues to make targeted, low-input marker systems a practical and necessary complement to genome-scale approaches.

The choice of genetic markers depends on both the research aims and the practical limitations of the study system (Andrews et al. 2016, Helyar et al. 2011). Genome-wide SNP datasets and whole-genome approaches are typically preferred for questions targeting broad evolutionary patterns or genome -scale variation. In contrast, studies that aim to differentiate and identify individuals, discriminate between closely related strains or relatives, illustrate recent demographic or dispersal events, or work with very small amounts of DNA may benefit from simpler marker systems. Microsatellites provide one such option. Their high allelic diversity and codominant inheritance allow accurate individual-level identification with relatively few loci, and their elevated mutation rates increase sensitivity to recent processes, including fine-scale demographic changes, replacement of genetic lineages, and natural or human-mediated shifts (Ellegren 2004, Vieira et al. 2016). At the same time, microsatellite panels require only modest laboratory effort and cost, which enables routine use, large sample sets, and processing that can be completed relatively quickly with standard local laboratory workflows (Hauser et al. 2021). Although they do not capture genome-wide patterns, microsatellites remain an efficient tool for targeted applications where individual discrimination, recent timescales, and operational simplicity are the main priorities.

Despite their practical advantages, developing robust microsatellites for *Gyrodactylus* has historically been challenging. Early work in this genus was conducted without genomic resources and relied on probe-enriched libraries from which only a limited number of cloned fragments could be screened, offering little certainty about the genomic context of candidate loci. In some cases, the material available for screening also offered only limited opportunity to detect polymorphism (Faria et al. 2011). Subsequent reference genomes have clarified why such approaches were difficult to apply. The *G. salaris* draft genome revealed substantial repetitive content and structurally complex regions that constrain the availability of unique, indel-free flanking sequences required for robust PCR amplification (Hahn et al. 2014), while comparative analyses of *G. bullatarudis* documented extensive gene duplication and paralogy that increase the risk of non-specific amplification, when primers are designed without genomic context (Konczal et al. 2020b). These genomic features highlight the value of genome-informed locus selection, in which putative microsatellites are evaluated *in silico* for motif structure, genomic uniqueness and flanking-sequence quality prior to laboratory validation.

Here, we present a *G. salaris*-specific microsatellite panel designed for individual-level genotyping. We use available genomic resources to identify candidate loci, compile these into a single-tube multiplex assay, and evaluate performance at the level of individual worms. The following sections outline the locus-selection workflow, primer design, multiplex optimization, and technical validation.

## Material and methods

### Marker development

Microsatellite candidates were first obtained through *in silico* repeat detection, using SciRoKo (Kofler et al. 2007), performed on preliminary genomic assemblies of *Gyrodactylus salaris* available to the authors for motif discovery. These assemblies were used solely as a search space for repetitive motifs and flanking regions. Minor base-level uncertainty, including potential homopolymer length variation, was expected in the design sequences and could affect predicted fragment length by a small margin. Expected amplicon sizes were therefore considered approximate.

#### Initial screening phase

From the repeat-based candidate set, an initial subset of loci was selected for empirical testing as a part of a broader pilot panel that also included SNPs and gene-based loci. This first screening phase, conducted in laboratory A, yielded three microsatellite loci with clear and reproducible amplification, while one additional locus was discarded due to inconsistent performance. The SNPs included in the pilot phase were not pursued further for the present study. Outcomes from this initial screening informed the refinement of the locus selection criteria for the expanded marker development.

#### Refined marker development

A more extensive and stringent *in silico* selection was subsequently performed using the same method on an updated preliminary assembly version available to the authors. Candidate loci were retained only if they met all these following mandatory requirements:

i. unique and sufficiently long flanking regions suitable for primer design,
ii. moderate repeat length (7–17 repeat units),
iii. localisation on different genomic fragments to minimise potential physical linkage, and
iv. depth of coverage consistent with single-copy regions in the draft assembly.

From the loci fulfilling these requirements, we selected a predefined set of candidates that were situated within introns or adjacent to annotated genes. These regions typically provide sufficiently unique flanking sequences and, together with the coverage information, increase the likelihood that the loci represent single-copy regions and allow their genomic positions to be cross-referenced with the chromosome-level reference assembly. Primer pairs for all refined candidates were designed in Primer3Plus using standard settings for GC content, melting temperature and amplicon length.

After completing the initial marker development work in laboratory A, the subsequent empirical steps were carried out in both laboratories. During a research visit to laboratory B, the existing screening-phase loci were first re-tested using laboratory-specific equipment and reagents. Additional refined-phase loci were then designed in laboratory B, together with preliminary multiplex configurations aimed at accommodating all available markers within a single multiplex format. New primer pairs were first verified in simple test reactions and then included in the multiplex evaluations. Loci that produced consistent fragment profiles across these iterative tests were returned to laboratory A for further optimization using the standard workflow of that laboratory. In total, twelve microsatellite loci met all technical requirements and were included in the final marker panel.

#### Mapping to the published Gyrodactylus genomes

To verify and report genomic locations of the final microsatellite loci, marker sequences together with their immediately available flanking regions were aligned post hoc to the published *G. salaris* scaffold-level reference assembly (Hahn et al. 2014). Each marker matched uniquely to a single reference sequence, allowing unambiguous placement at the scaffold or contig level. Genomic coordinates are reported using reference identifiers and positional coordinates relative to this assembly (Table 3).

To obtain chromosome-level context in a second species, marker sequences were extended with up to 2 kb of flanking sequence on each side, using available sequence only, and aligned to the chromosome-level *G. kobayashii* reference genome (Zhang et al. 2025) using nucleotide-level sequence similarity. Resulting matches were used to assign the chromosome on which the corresponding genomic region is located. These assignments provide contextual information only and do not imply precise localization or chromosomal organization of the loci in *G. salaris*.

### PCR conditions and multiplex configuration

All forward primers were tagged with different fluorescent labels, and the resulting amplicons spanned approximately 150–600 bp. This size distribution allowed all loci under evaluation at that stage to be accommodated within a single multiplex reaction.

All loci under evaluation were amplified in the same reaction using the Type-it Microsatellite PCR kit (Qiagen). The amplification reaction was done in a 6 µl volume and it consisted of 1 µl of template DNA, 1x Type-it Multiplex PCR Master Mix (Qiagen), and 0.2 µM of each primer, except for loci 697 and 21946 0.4 µM of F primer and 0.2 µM of each R primer, and sterile H_2_0. The amplification method was as follows: 95 °C for 5 min, 30 cycles of 95 °C for 30 s, 60 °C for 90 s and 72 °C for 30 s, the final extension in 68 °C for 30 min and cooling down to 4 °C. The PCR products were checked on a 1.5 % agarose gel.

For some loci with longer expected amplicons, additional primer pairs were designed to produce shorter fragments to improve amplification consistency. The shorter versions of loci 22089, 25338, 5780 and 21946B were amplified either in a separate four-locus reaction or multiplexed with other microsatellite markers, depending on the assay configuration.

To make sure that amplification was specific, microsatellite alleles were also confirmed by sequencing. The amplification was done for each locus separately. The reaction consisted of 1x Phusion buffer (Finnzymes), mM of each dNTP, 1μM of each primer, 0.2U of Phusion DNA polymerase, and sterile H_2_0. The amplification method was as follows: 98 °C for 1 min, 35 cycles of 98 °C for 10 s, 60 °C for 20 s and 72 °C for 1 min, the final extension in 72 °C for 10 min and cooling down to 4 °C. The PCR products were checked on a 1.5 % agarose gel and purified using the Exo-FastAP method (Fermentas). The purification reaction was done in 10 μl volume and it consisted of 2 μl of PCR product, 0.8x FastAP buffer, 0.5U FastAP, 1U ExoI, and sterile H_2_0. The sequencing reactions were performed using the BigDye Terminator v3.1 Cycle Sequencing Kit according to manufacturer’s instructions.

### Genotyping and allele scoring

Fragment analysis was performed on an ABI 3730 sequencer according to the manufacturer’s instructions. Fragment sizes were determined using a manufacturer-supplied internal size standard appropriate for the instrument. GeneMapper v4.1 (Applied Biosystems) was used to analyze DNA fragments and to score genotypes. Alleles were called based on consistent fragment profiles, and all genotype calls were manually reviewed to ensure scoring consistency across loci. Fragment sizes were assigned to predefined allele bins with a ±1 bp tolerance. For loci where shorter amplicon versions were used, the resulting fragment sizes were converted to the corresponding size classes of the original full-length amplicons to ensure consistency in allele scoring.

To verify locus specificity and to check for potential indels outside the repeat region that could influence fragment length, a small subset of amplicons was sequenced using standard Sanger methods. Sequence chromatograms were visualized and edited in CodonCode Aligner v9.0.1 (CodonCode Corporation) and MEGA X (Kumar et al. 2018), and manual corrections were applied where required to ensure accurate base calling.

### Validation material

#### Studied G. salaris specimens

A large pool of individual *G. salaris* specimens (approximately 1600 individuals) was available for DNA template preparation and for evaluating the performance of the microsatellite assays. The specimen set originated from routine diagnostics, regulatory surveillance and a variety of research sampling activities conducted by multiple organisations. Specimens were preserved in 96 % ethanol either individually or as several individuals stored together by sampling occasion. The material covered a wide geographic range of the species, providing diverse template conditions for methodological validation. This allowed us to assess amplification success, fragment profile consistency and multiplex robustness across diverse templates. For analyses that required higher-quality template DNA, an internal subset of approximately 800 specimens was used. This subset was selected based on DNA quality rather than biological criteria.

#### DNA template preparation

For molecular analysis, single parasite specimens were picked from the ethanol-preserved storage tubes, irrespective of whether the tubes contained individual specimens or several specimens collected during the same sampling occasion. Prior to enzymatic DNA release, the ethanol was evaporated from each specimen.

DNA templates were prepared using three laboratory-specific protocols, reflecting differences in equipment and routine workflows between laboratories rather than biological considerations:

##### 1. Proteinase K lysis followed by DNA extraction (GeneMole)

Specimens were lysed in 100 µl of lysis buffer and 5 µl of proteinase K, and 100 µl of the lysate was used for DNA extraction with the GeneMole instrument (Mole Genetics AS) following the manufacturer’s DNA tissue protocol.

##### 2. PrepMan® Ultra DNA release

For some specimens, DNA was released using PrepMan® Ultra (Invitrogen). After ethanol evaporation, 50 µl of PrepMan buffer was added, followed by incubation for 15 min at 94 °C and cooling to 4 °C.

##### 3. Proteinase K lysis without purification (direct-lysate template)

DNA was released using a proteinase K lysis protocol and lysates were used directly as PCR templates without further purification. The lysis program consisted of incubation at 65 °C for 25 min, followed by 94 °C for 10 min to inactivate proteinase K, and cooling to 4 °C.

All approaches yielded PCR-ready DNA templates. After DNA release, lysates or extracts were stored at −20 °C until use.

#### Cross-species amplification

Cross-species amplification was evaluated using a broad panel of non-target *Gyrodactylus* specimens representing more than 20 nominal species and lineages. All reactions were performed under the same PCR conditions as used for *G. salaris*, without species-specific optimisation.

## Results

A total of twelve microsatellite loci met all technical criteria during the two-phase development process and were included in the final marker panel, representing twelve distinct genomic regions. Two of the loci contained compound repeat structures with two adjacent repeat motifs each, resulting in fourteen repeat blocks in total. Primer sequences, fluorescent labels, repeat motifs, and expected amplicon sizes are summarized in Table 1. Technical performance metrics, including the number of detected alleles, allelic size ranges and amplification success rates, are presented in Table 2. Genomic positions inferred from the *G. salaris* draft assembly and sequence similarity based chromosomal assignments in *G. kobayashii* are provided separately in Table 3.

**Table 1.**
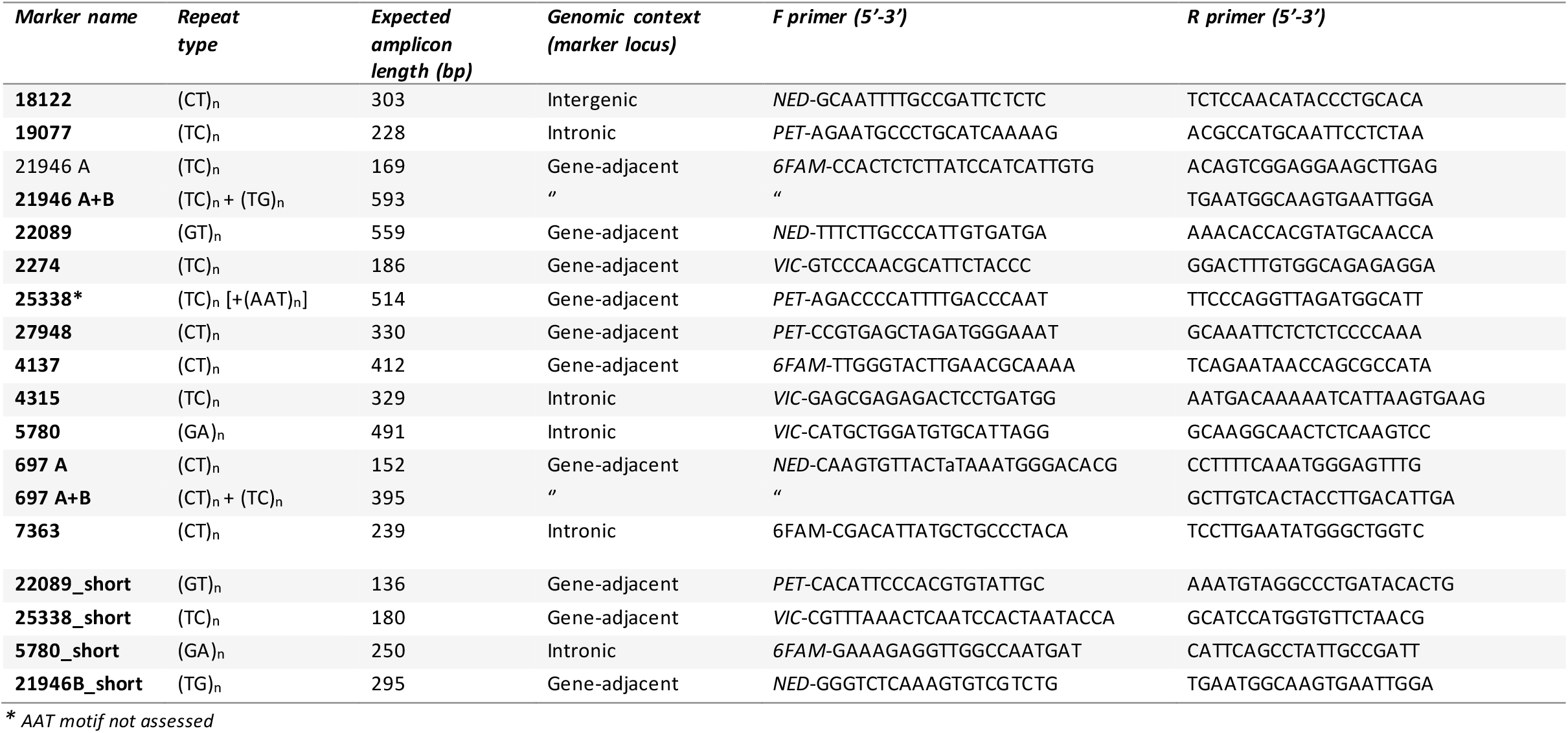
Designed microsatellite markers for this study. Naming of loci, repeat structures, and expected product lengths is based on the in silico primer design stage using 208 reference sequence information available to the authors. Genomic context assignments are based on preliminary internal annotation. Intronic refers to physical overlap of 209 the marker locus with an annotated intron, regardless of strand orientation. The forward primers were labelled with four fluorescent dyes (6FAM, VIC, NED and PET). Two 210 loci (697 and 21946) contained two adjacent repeat blocks (A and B). These compound loci were genotyped using one labelled forward primer to gether with two reverse 211 primers. Repeat motifs are reported in the forward primer orientation, which defines the operational orientation of the marker.

**Table 2.** Technical characterization of microsatellite loci. Allele numbers, fragment-size ranges, and amplification success are shown for the full dataset (“Full”, N = 1603) 235 and the high-quality subset (“HQ”, N = 809). Size range is reported for the long amplicon versions used in the original multiplex design. Short amplicon primer pairs, whe re 236 available, are listed in Table 1 and were not used for size range estimation.

| Locus | Allele count<br>(observed allele classes) |  | Size range (bp) |  | Amplification success (%) |  | Notes |
| --- | --- | --- | --- | --- | --- | --- | --- |
|  | Full | HQ | Full | HQ | Full | HQ |  |
| **18122** | 9 | 7 | 282-302 | 286-302 | 95,8 | 100 |  |
| **19077** | 7 | 4 | 216-228 | 222-228 | 98,3 | 100 |  |
| 21946A* | 8 | 7 | 152-166 | 158-172 | 99,5 | 100 |  |
| **21946 (A+B)** | 9 | 9 | 582-602 | 582-602 | 86,5 | 100 | An additional variable short indel outside the repeat |
| **22089** | 10 | 10 | 552-576 | 552-576 | 92,4 | 100 | An additional variable indel outside the repeat region<br>contributes to length variation in the longer amplicon but is<br>not present in the short amplicon. |
| **2274** | 9 | 7 | 172-188 | 174-188 | 99,3 | 100 |  |
| **25338\*\*** | 44 | 42 | 492-586 | 496-586 | 72,5 | 100 | Very long repeats may show technical length variation |
| **27948** | 11 | 9 | 314-336 | 314-334 | 95,5 | 100 |  |
| **4137** | 8 | 7 | 392-414 | 392-412 | 92,5 | 100 |  |
| **4315** | 22 | 16 | 304-362 | 318-362 | 95,6 | 100 |  |
| **5780** | 19 | 16 | 460-512 | 466-512 | 93,1 | 100 |  |
| 697A* | 33 | 32 | 137-207 | 137-207 | 97,9 | 100 | Very long repeats may show technical length variation |
| **697 (A+B)** | 51 | 45 | 235-449 | 237-447 | 84,2 | 100 | Additional variable indels outside the repeat:<br>(i) a long indel between the A and B repeat<br>(ii) a short indel between B repeat and the R primer |
| **7363** | 42 | 35 | 215-297 | 223-289 | 88,0 | 100 | Very long repeats may show technical length variation |

\* A = the first repeat array only; **A+B** = full compound repeat at the same genomic locus.
\*\* Only (CT)<sub>n</sub> alleles scored; flanking AAT motif not assessed.

**Table 3.** Genomic placement of the developed markers. Markers were mapped to the scaffold-level reference genome of *G. salaris* (Hahn *et al*. 2014). Structural variation observed in the study material is reported separately (Table 2) and is not reflected in the reference-based annotation shown here. Orthologous regions were verified by BLASTn searches against the chromosome-level genome of *G. kobayashii* (Zhang *et al*. 2025).

| Marker | Reference position in *G. salaris*<br>(*Gsalaris_v1*, *GCA_000715275.1*; *PRJNA244375* - *Lier*) | Strand | Chromosomal position in *G. kobayashii*<br>(*GWHFQJJ000000000.1*) |
| --- | --- | --- | --- |
| 18122 | contig_scf7180006953077_0: 38,590–38,894 | plus | no reliable placement |
| 19077 | contig_scf7180006952609_0: 29,170–28,948 | minus | Chr4: ~5,3 Mb (GWHFQJJ000000004.1) |
| 21946 | contig_scf7180006949357_0: 23,758–24,352 | plus | Chr3: ~8,0 Mb (GWHFQJJ000000003.1) |
| 21946B_short | contig_scf7180006949357_0: 24,052–24,353 | " | " |
| 22089 | contig_scf7180006953358_2: 10,786–10,215 | minus | no reliable placement |
| 22089_short | contig_scf7180006953358_2: 10,540–10,405 | " | " |
| 2274 | contig_scf7180006948928_2: 12,985–13,171 | plus | Chr1: ~10,4 Mb (GWHFQJJ000000001.1) |
| 25338 | contig_scf7180006950910_0: 890–1,380 | plus | no reliable placement |
| 25338_short | contig_scf7180006950910_0: 1,096–1,255 | " | " |
| 27948 | contig_scf7180006948470_2: 18,943–18,629 | minus | no reliable placement |
| 4137 | contig_scf7180006950433_0: 3,465–3,870 | plus | no reliable placement |
| 4315 | contig_scf7180006950445_0: 1,399–1,077 | minus | no reliable placement |
| 5780 | contig_scf7180006953409_1: 40,307–39,781 | minus | Chr2: ~4,0 Mb (GWHFQJJ000000002.1) |
| 5780_short | contig_scf7180006953409_1: 40,183–39,898 | " | " |
| 697 | contig_scf7180006948120_0: 40,031–40,431 | plus | no |
| 7363 | contig_scf7180006953439_0: 31,452–31,690 | plus | Chr5: ~1,8 Mb (GWHFQJJ000000005.1) |

### Primer information

Table 1 summarizes the primer design information for all twelve loci, including locus names, repeat structures, expected product lengths, and primer sequences with their fluorescent labels. These parameters derive from the *in silico* design stage and describe the intended amplification products prior to empirical testing. All loci were designed to be amplified under a single multiplex PCR protocol. Two loci (697 and 21946) contained two adjacent repeat blocks (A and B). For these compound loci, one labelled forward primer was paired with two reverse primers targeting the respective repeat regions. The genomic context column provides a brief positional description of each amplicon based on available design-phase sequence information and does not imply any functional interpretation. Markers were classified as intronic, gene-adjacent, or intergenic based on design-phase criteria.

### Locus characterization

Table 2 summarizes the empirical performance of the twelve loci, including allele counts, observed size ranges, and amplification success rates. Only specimens that yielded a PCR product for at least one locus were included in the reported datasets. All loci were polymorphic and produced scorable fragment profiles under the optimized PCR conditions. Amplification success was consistently high, and allele profiles were stable across replicate PCRs. The reported metrics describe the technical performance of the loci and are not intended to imply population-level or biological interpretation.

Amplification percentages in Table 2 reflect successful reactions from both the primary full-length assays and, for some loci, additional primer sets targeting shorter amplicons. These primers were used only when the long amplicon of the primary assay failed to amplify due to low template quantity or DNA degradation. Their inclusion increases the overall amplification percentage for the affected loci, because samples that would otherwise score as failures under the long-amplicon assay are recovered using the shorter fragment.

Independently of amplicon length, loci with long repeat arrays (e.g. 25338) showed a greater number of observed alleles and a wider size range in the full dataset. This reflects the combined effects of true biological length variation associated with long, structurally heterogeneous repeat tracts, and the broader technical size dispersion that occurs when alleles differ by many repeat units. In the high-quality subset these same loci display slightly fewer alleles and a narrower size range, indicating that the allele surplus in the full dataset arises from both biological heterogeneity of the long sequence and the technical resolution limits associated with long fragments.

Across all loci, allele scoring remained unambiguous in the high-quality dataset, indicating that the observed variation did not compromise the reliability of the marker panel.

### Additional notes on locus performance

All twelve loci produced clear and scorable fragment profiles under the optimized amplification conditions. Allele calls were consistent across replicate PCRs, and no locus showed evidence of allele dropout or nonspecific products.

One locus (7363) displayed slightly more variable peak intensities across replicates, but the signal remained sufficient for unambiguous allele categorization. For loci with very long repeat arrays, fragment size variation occasionally exceeded single-base resolution, but alleles nevertheless formed clearly separable size classes, and categorization remained consistent when considering the full peak pattern together with supporting information from the other loci.

In high-quality templates, nearly all loci amplified robustly in the multiplex reaction. The longest amplicons, however, showed dependable amplification only in samples with intact and adequate DNA, consistent with their larger fragment size. All other loci produced tightly clustered peak profiles across runs, and no cross-amplification was detected in negative controls. Taken together, these results indicate that the marker panel performs reliably across individuals under typical single-worm DNA yields, with only the longest amplicons showing sensitivity to template quality. This demonstrates that the multiplex can be applied effectively under the DNA constraints inherent to single-worm extractions.

When aligning the full 25338 amplicon sequence to the published *G. salaris* genome, a short AAT repeat located at the margin of the amplicon was found to differ by one repeat unit from the internal reference sequence, indicating that this flanking motif may be polymorphic. In the current multiplex configuration, the locus was designed and scored based on variation in the central (CT)_n_ repeat, with the reverse primer annealing immediately adjacent to the (AAT)_n_ sequence (5′-AAT-3’). With this primer placement, potential length variation in the flanking (AAT)_n_ motif is substantially smaller than the variation in the (CT)_n_ array and cannot be resolved separately in fragment analysis. Accordingly, only (CT) _n_ alleles were analyzed in this study.

Primer designs enabling amplification of a shorter fragment that excludes the flanking (AAT)_n_ motif were also generated and are reported for this locus (25338_short). However, the potential analytical implications of targeting shorter versus longer amplicons were not examined systematically in this study. A detailed assessment of variation in the adjacent (AAT)_n_ repeat, or of alternative primer configurations separating the (AAT)_n_ and (CT)_n_ motifs, would therefore require dedicated analysis beyond the scope of the present work.

Two loci (21946 and 697) contain two adjacent repeat modules (“A” and “B”), forming a compound repeat structure. In the multiplex PCR, the full A+B region was amplified using a labelled forward primer together swith a reverse primer located downstream of the B repeat, producing a long compound amplicon. A second reverse primer, positioned immediately downstream of the A repeat, yielded a shorter A only amplicon, which can be used to obtain an independent estimate of the A repeat length.

Sequence inspection of the extended 697 locus sequence showed that, in addition to variation within the A and B repeat modules, length differences may occur (i) in the short intervening segment between the two repeat modules and (ii) in the region immediately upstream of the reverse primer binding site (Table 2). These indels contribute to the overall length of the long A+B amplicon and therefore prevent reliable derivation of the B repeat variation from the compound fragment alone. Accurate B repeat estimates require joint analysis of the A only and A+B amplicons. The B repeat cannot be isolated directly from the compound product.

In this study, routine genotyping relied solely on the combined 697 A+B amplicon, and B only repeat counts were not derived. The long A+B amplicon produced clear and reproducible peak profiles across the validation material, and the presence of intervening indels did not affect the formation of distinct size classes or allele categorization. The shorter A only amplicon amplified readily and may be useful in applications where only the first repeat module is required.

If an amplicon displays unexpected or ambiguous fragment length behavior, confirmation by sequencing is recommended, as this allows discrimination between variation within the repeat modules and indels occurring between or adjacent to them.

### Genomic placement

All loci were aligned post hoc to the published *G. salaris* scaffold-level reference assembly (Hahn et al. 2014). Each locus mapped uniquely to a single reference sequence, allowing unambiguous reporting of genomic positions at the scaffold level. The corresponding reference identifiers and genomic positions are summarized in Table 3.

For some loci, sequence similarity searches returned matches to both an unplaced scaffold and a contig within the same reference assembly. Inspection of the corresponding genomic regions showed that these entries represent the same underlying sequence rather than independent loci, reflecting assembly structure rather than biological duplication. In such cases, contig-based coordinates were used for reporting genomic positions. Strand orientation (+/−) refers to the orientation relative to the reference genome assembly.

To provide chromosome-level context in a second species, loci were associated with the chromosome-level genome assembly of *G. kobayashii* (Zhang et al. 2025). Chromosomal assignment was inferred based on nucleotide-level sequence similarity using the amplicon sequence extended with up to 2 kb of flanking sequence on each side. These assignments indicate the chromosome on which the corresponding genomic region is located and are reported for contextual purposes only. They do not imply precise localization or physical linkage of the microsatellite loci in *G. salaris*.

### Cross-species amplification

Amplification products were obtained from some nontarget specimens, but this was not consistent across loci or specimens. Cross-amplification alone cannot be interpreted as evidence of marker transferability. Because the markers target noncoding regions, amplification outside *G. salaris* cannot be assumed to represent orthologous loci without sequence confirmation. The marker panel is therefore intended for use in studies of *G. salaris*, and application beyond this context requires independent validation.

## Discussion

The twelve microsatellite loci developed in this study provide a reliable way to obtain nuclear, individual-level genetic data for a species where each organism yields very little DNA. All loci amplified consistently under the optimized PCR conditions, produced clear and scorable fragment profiles, and were polymorphic at the allele level. Minor variation in fragment-size distributions for the longest amplicons was accommodated with slightly broader allele-binning thresholds, in line with patterns reported for long microsatellite products in methodological studies (Ellegren 2004, Vieira et al. 2016). These features did not affect replicate consistency or allele calling, demonstrating that the method yields reproducible, fragment-based genotypes from single worms.

All loci amplified reliably in a single multiplex PCR. Running all loci in one multiplex reaction is particularly advantageous for species where each individual produces only very small quantities of DNA. A single *Gyrodactylus* individual contains on the order of ∼1000 cells (Bakke et al. 2007) and yields approximately 5 ng of high-molecular-weight genomic DNA under standard proteinase K lysis conditions (Cunningham & Mo 1997), making efficient use of template essential. This multiplex setup therefore enables rapid and cost-effective processing while requiring only minimal amounts of DNA. The combination of consistent amplification and informative allele-level patterns across loci provides the level of individual discrimination needed for a wide range of genetic applications, including comparative analyses across sampling events, assessments of genetic relatedness, and other studies that benefit from repeatable nuclear data at the scale of single parasites. These same properties also facilitate practical tasks such as routine genotyping within monitoring programs or confirming repeated detections in surveillance contexts. When whole-genome approaches require pooling or are otherwise impractical due to template limitations, multiplexed microsatellites provide a scalable and efficient alternative for generating repeatable nuclear data from single parasites.

In addition to microsatellite repeat length variation, indel polymorphism outside the repeat region was observed in several PCR amplicons during marker validation. This variation was not a target of marker design and became apparent only through downstream analyses. While such indels can complicate fragment-based genotyping when combined with repeat length variation, they may provide biologically informative signals in sequence-based analyses, for example in studies addressing population structure or dispersal history. Importantly, the ability to derive both fragment-based and sequence-based information from the same PCR products highlights a practical advantage of a compact and well-characterized marker set, allowing flexible analytical strategies depending on the research question.

Genome-informed locus selection was an important consideration in assembling a technically robust microsatellite panel for this taxonomic group. Although no previous studies have developed variable microsatellite markers for *G. salaris*, available *Gyrodactylus* genome resources indicate structural features that complicate purely de novo marker development. High repeat content and locally heterogeneous sequence structure have been documented in the *G. salaris* draft genome (Hahn et al. 2014), and similar features have been reported in related species (Konczal et al. 2020b). In such genomic backgrounds, the uniqueness and stability of primer binding regions cannot be assumed a priori, increasing the risk of unintentional co-amplification of duplicated or closely related loci if marker development is performed without reference sequence information.

Genome-informed locus selection can help evaluate primer binding sites in relation to local sequence structure, reducing the likelihood of unintended co-amplification and increasing confidence in the structural independence of the resulting loci. While the present in silico analyses do not resolve chromosomal positions for the loci in *G. salaris*, the ability to anchor a subset of markers to published genome assemblies demonstrates practical applicability for future integration as higher-resolution genomic resources become available. Definitive chromosomal assignments would require either a chromosome-level reference genome for *G. salaris* or independent cytogenetic validation.

Although microsatellites do not substitute for genome-wide datasets or capture adaptive variation (Andrews et al. 2016, Hoban 2016), they remain an efficient option for targeted applications that require individual-level resolution on recent timescales and must operate under stringent DNA constraints. Within this scope, the marker set developed here offers a technically robust, multiplex-compatible tool for routine fragment-based genotyping of *G. salaris*. The core methodological outcome of this work is the demonstration that nuclear genotyping at the level of single worms is technically achievable in this system, with sufficient allele-level resolution to generate consistent and repeatable data under standard laboratory conditions.

## Data accessibility

All data supporting the results of this study are provided in the manuscript.

## Authors’ contributions (based on CRediT)

Conceptualization: HA, HH & JL; Methodology: HA; Validation: HA; Investigation: HA; Resources: HH, JL; Data curation (including specimen curation): HA, HH, JL; Writing – original draft: HA; Writing – review & editing: HA, HH & JL

## Acknowledgements

We acknowledge the contribution of all individuals and organizations involved in sample collection. We thank Laura Törmälä for extensive laboratory work during the marker validation phase, and Soile Alatalo and Hannele Parkkinen for generating the sequencing and fragment analysis data used in this study.

## References

Allendorf, F.W., Hohenlohe, P.A. & Luikart, G. (2010). Genomics and the future of conservation genetics. Nature Reviews Genetics, 11, 697–709.

Andrews, K.R., Good, J.M., Miller, M.R., Luikart, G. & Hohenlohe, P.A. (2016). Harnessing the power of RADseq for ecological and evolutionary genomics. Nature Reviews Genetics, 17, 81–92.

Bakke, T.A., Cable, J. & Harris, P.D. (2007). The biology of gyrodactylid monogeneans: the “Russian-doll killers”. Advances in Parasitology, 64, 161–376.

Cunningham, C.O. & Mo, T.A. (1997). Random amplified polymorphic DNA (RAPD) analysis of three Norwegian Gyrodactylus salaris populations (Monogenea: Gyrodactylidae). Journal of Parasitology, 83(2), 311–314.

Ellegren, H. (2004). Microsatellites: simple sequences with complex evolution. Nature Reviews Genetics, 5, 435–445.

Faria, P.J., Harris, P.D., Julca, I., Bakke, T.A. & Bachmann, L. (2011). First polymorphic microsatellites for the gyrodactylids (Monogenea), an important group of fish pathogens. Conservation Genetics Resources, 3, 177–180.

Hahn, C., Fromm, B. & Bachmann, L. (2014). Comparative genomics of flatworms (Platyhelminthes) reveals shared genomic features of ecto- and endoparasitic Neodermata. Genome Biology and Evolution, 6, 1105–1117.

Hansen, H., Bakke, T.A. & Bachmann, L. (2007). DNA taxonomy and barcoding of monogenean parasites: lessons from Gyrodactylus. Trends in Parasitology, 23, 363–367.

Hauser, S.S., Athrey, G. & Leberg, P.L. (2021). Waste not, want not: Microsatellites remain an economical and informative technology for conservation genetics. Ecology and Evolution, 11, 15800–15814.

Helyar, S.J., Hemmer-Hansen, J., Bekkevold, D., Taylor, M.I., Ogden, R., Limborg, M.T., Cariani, A., Maes, G.E., Diopere, E., Carvalho, G.R. & Nielsen, E.E. (2011). Application of SNPs for population genetics of non-model organisms: new opportunities and challenges. Molecular Ecology Resources, 11, 123–136.

Hodel, R.G.J., Segovia-Salcedo, M.C., Landis, J.B., Crowl, A.A., Sun, M., Liu, X., Gitzendanner, M.A., Douglas, N.A., Germain-Aubrey, C.C., Chen, S., Soltis, D.E. & Soltis, P.S. (2016). The report of my death was an exaggeration: a review for researchers using microsatellites in the 21st century. Applications in Plant Sciences, 4(6), 1600025.

Hohenlohe, P.A., Funk, W.C. & Rajora, O.P. (2021). Population genomics for wildlife conservation and management. Molecular Ecology, 30(1), 62–82.

Kofler, R., Schlötterer, C. & Lelley, T. (2007). SciRoKo: a new tool for whole genome microsatellite search and investigation. Bioinformatics, 23, 1683–1685.

Konczal, M., Przesmycka, K.J., Mohammed, R.S., Phillips, K.P., Camara, F., Chmielewski, S., Hahn, C., Guigó, R., Cable, J. & Radwan, J. (2020a). Gene duplications, divergence and recombination shape adaptive evolution of the fish ectoparasite Gyrodactylus bullatarudis. Molecular Ecology, 29, 1494–1507.

Konczal, M., Przesmycka, K.J., Mohammed, R.S., Hahn, C., Cable, J. & Radwan, J. (2020b). Expansion of frozen hybrids in the guppy ectoparasite Gyrodactylus turnbulli. Molecular Ecology, 30, 1005–1016.

Kumar, S., Stecher, G., Li, M., Knyaz, C. & Tamura, K. (2018). MEGA X: Molecular Evolutionary Genetics Analysis across Computing Platforms. Molecular Biology and Evolution, 35, 1547–1549.

Luikart, G., England, P.R., Tallmon, D., Jordan, S. & Taberlet, P. (2003). The power and promise of population genomics: from genotyping to genome sequencing. Nature Reviews Genetics, 4, 981–994.

Vieira, M.L.C., Santini, L., Diniz, A.L. & Munhoz, C.F. (2016). Microsatellites: what they mean and why they are so useful. Genetics and Molecular Biology, 39, 312–328.

Zhang, D., Zhao, J-M., Xiang, C-Y., Ma, Y-W., Lei, H-P., Shi, Y-Y., Zhou, S., Zeng, X., Chen, J., Liu, F., Zeng, B., Song, R., Hu, Y., Zhang, F., Liu, X., Li, W.-X.Wang, G.-T. & Jakovlić, I. (2025). Genome-wide protein interaction analysis in parasitic Gyrodactylus flatworms–fish host systems and drug target identification. Advanced Science, 12, e14618.

